# Imperfect drug penetration leads to spatial monotherapy and rapid evolution of multi-drug resistance

**DOI:** 10.1101/013003

**Authors:** Stefany Moreno-Gamez, Alison L. Hill, Daniel I. S. Rosenbloom, Dmitri A. Petrov, Martin A. Nowak, Pleuni Pennings

**Author notes:** These authors contributed equally to the manuscript (Stefany Moreno-Gamez, Alison L. Hill).

## Abstract

Infections with rapidly evolving pathogens are often treated using combinations of drugs with different mechanisms of action. One of the major goals of combination therapy is to reduce the risk of drug resistance emerging during a patient’s treatment. While this strategy generally has significant benefits over monotherapy, it may also select for multi-drug resistant strains, which present an important clinical and public health problem. For many antimicrobial treatment regimes, individual drugs have imperfect penetration throughout the body, so there may be regions where only one drug reaches an effective concentration. Here we propose that mismatched drug coverage can greatly speed up the evolution of multi-drug resistance by allowing mutations to accumulate in a stepwise fashion. We develop a mathematical model of within-host pathogen evolution under spatially heterogeneous drug coverage and demonstrate that even very small single-drug compartments lead to dramatically higher resistance risk. We find that it is often better to use drug combinations with matched penetration profiles, although there may be a trade-off between preventing eventual treatment failure due to resistance in this way, and temporarily reducing pathogen levels systemically. Our results show that drugs with the most extensive distribution are likely to be the most vulnerable to resistance. We conclude that optimal combination treatments should be designed to prevent this *spatial effective monotherapy*. These results are widely applicable to diverse microbial infections including viruses, bacteria and parasites.

## 1 Introduction

Current standard-of-care treatment for many bacterial and viral infections involves combinations of two or more drugs with unique mechanisms of action. There are two main situations in which *combination therapy* significantly outperforms *monotherapy* (treatment with a single drug). Firstly, in clinical scenarios where precise pathogen identification is not possible before treatment begins (“empirical therapy”), or when infections are suspected to be polymicrobial, treating with multiple drugs increases the chances of targeting the virulent organism. Secondly, even when infections are caused by a single, precisely identified microbe, combination therapy reduces the risk of developing drug resistance. This reduced risk is believed to follow from the fact that multiple mutations are generally needed to enable pathogen growth when multiple drugs are present. In addition, the use of multiple drugs may reduce the residual population size and thus further reduce the rate of evolution of resistance. Preventing the evolution of resistance is particularly relevant to infections caused by rapidly evolving pathogens, and to persistent infections that can be controlled but not cured, for which there may be a high risk of drug resistance evolving during the course of a single patient’s treatment. Despite wide-spread use of combination therapy, drug resistance remains a serious concern for many infections in this category, such as HIV, HBV, HCV and TB (1–4), as well as for certain cancers (5, 6). Understanding the factors that facilitate the evolution of multi-drug resistance is therefore a research priority.

Combination therapy can be compromised by treatment regimes that allow resistance mutations to different drugs to be acquired progressively (i.e., in stepwise fashion) rather than concurrently. This can occur when only one drug out of the combination is active during certain time periods. For example, starting patients on a single drug before adding a second drug promotes the evolution of multi-drug resistance(7, 8). A similar effect is seen for studies that rotate antibiotics (9–11). Even if drugs are given simultaneously but have different *in vivo* half-lives, periods of “effective monotherapy” with the longer-lived drug can occur, which may favor resistance evolution (12–14). HIV is one pathogen for which evolution of drug resistance is well studied. Surprisingly, it has been found that stepwise evolution of drug resistance is common in treated HIV^+^ individuals (15–17). It is unclear whether periods of effective monotherapy can explain this observation.

While many recent studies have focused on the potential impact of different half-lives between drugs, much less is known about how the spatial distribution of drugs influences the evolution of multi-drug resistance during combination therapy. Many treatments may involve mismatched drug penetrability – that is, there may be regions of the body where only a subset of drugs within a combination reach a therapeutic level (18). For example, many anti-HIV drugs have been observed at subclinical concentrations in the central nervous system and the genital tract (19, 20). Low concentrations in these body compartments, even when plasma concentrations are high, may allow viral replication and selection of resistance mutations (18, 21, 22), which may eventually migrate to the blood and lead to treatment failure (23). In another example, poor antibiotic penetration within biofilms (24) or certain body tissues (25) during treatment for *Staphylococcus aureus* infections is again associated with resistance evolution. Some recommend that this problem be addressed by pairing drugs with high efficacy but low penetration with other drugs of higher penetration, so that total drug coverage in the body increases (25). Yet this is likely a risky strategy. We hypothesize that combination therapy with drugs that have different penetration profiles will generally be more vulnerable to resistance, as it promotes situations of effective monotherapy that may allow a migrating pathogen lineage to acquire resistance mutations in a stepwise manner.

Previous work on the effect of drug penetration on drug resistance has focused on monotherapy. A mathematical model of viral infections showed that the window of drug concentrations where resistance mutations can arise and fix is greatly increased if there is a “drug-protected compartment” or “drug sanctuary” – a place where the drug level is not high enough to prevent virus replication (26). More recent theoretical work has explored the role of concentration gradients in the evolution of antibiotic resistance. This work demonstrated that when multiple mutations are needed for resistance to a single drug, either a continuous concentration gradient (27) or discrete microenvironments with differing concentrations (28) can speed up the rate of evolution. Experiments in microfluidic chambers where mobile bacteria grow in the presence of a spatial drug concentration gradient have confirmed that adaptation is accelerated (29). These results are surprisingly similar to studies that create temporal gradients in drug concentrations (30).

In this paper, we examine the role that drug penetration plays in evolution of resistance during combination therapy – thereby addressing a broad range of effective drug treatments. Specifically, we use a mathematical modeling strategy to show how the existence of anatomical compartments where only single drugs are present can drastically change the rate at which multi-drug resistance emerges. Among several pharmacologic and genetic determinants of resistance, we find that the size of single-drug compartments is key. A simple mathematical expression describes the critical size of single-drug compartments above which drug resistance emerges at an elevated rate, due to stepwise accumulation of mutations. In addition, we discover that combination therapy strategies face a general trade-off between suppressing microbial growth throughout the entire body and preventing eventual emergence of multidrug resistance. This trade-off implies, perhaps counterintuitively, that it may be rational to allow low-level microbial growth restricted to a small compartment where no drugs penetrate, in order to avoid regions of mismatched drug penetration – and increased risk of resistance emerging in the entire body. We discuss implications of this work for designing optimal drug combinations to prevent spatial effective monotherapy. Finally, we use our theory to explain why stepwise evolution of resistance may occur during effective combination therapy, as is sometimes seen clinically.

## 2 Model

Our goal is to understand the role of drug penetration in the evolution of multi-drug resistance. We consider an individual patient’s body to be divided into discrete and interconnected compartments where each drug either effectively suppresses pathogen growth or is completely absent (Figure 1). We model microbial dynamics in this environment, including growth, mutation, competition between strains, and migration between compartments. For simplicity, we focus on the case of two drugs only, though extensions to combinations of three or more drugs are straight-forward

**Figure 1:**
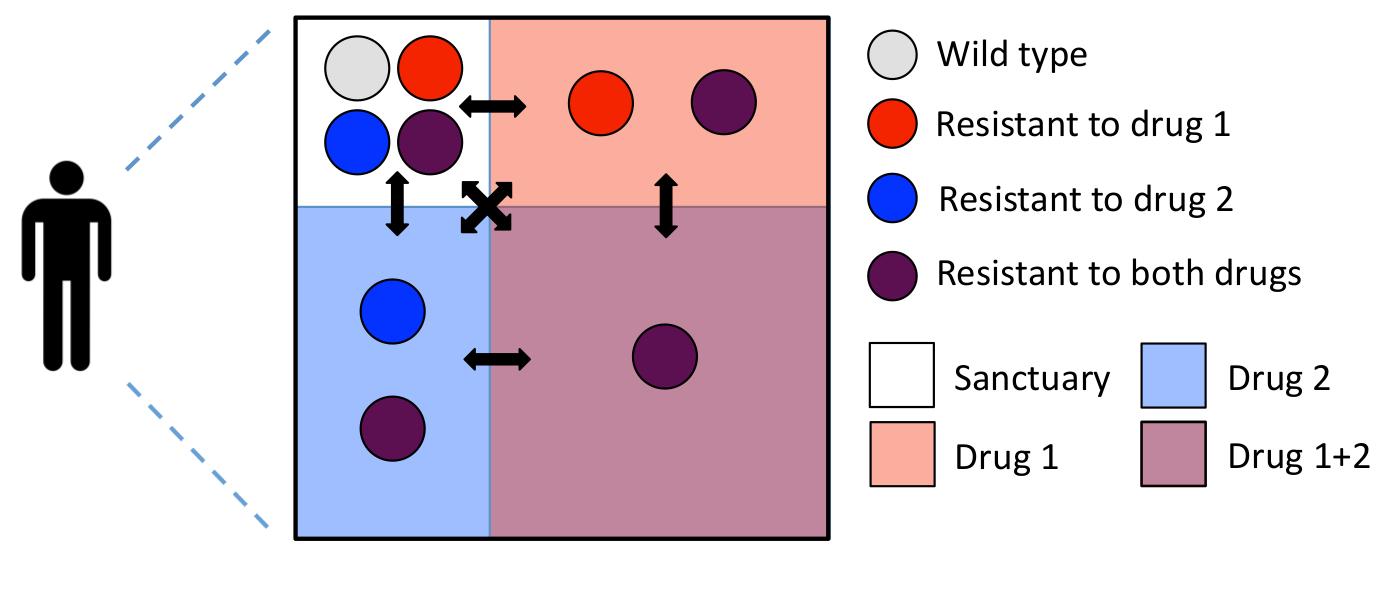
Compartment model for combination therapy with two drugs. The box represents a patient’s body and the red and blue shaded areas indicate the presence of drug 1 and 2 respectively. Mismatched drug penetration creates regions in the body where only one drug from the combination is present. We refer to these regions as single-drug compartments. Colored circles represent the pathogen genotypes: wild type (light gray), mutant resistant to drug 1 (red), mutant resistant to drug 2 (blue) and double-drug resistant mutant (purple). In the sanctuary all the pathogen genotypes can grow since none of the drugs is present. In the single-drug compartments only pathogens carrying a resistance mutation against the active drug can grow; that is, each drug alone suppresses pathogen growth. Finally, in the double-drug compartment only the double-drug resistant mutant can grow. All the compartments are connected by migration as indicated by the black arrows. Treatment fails when the double-drug compartment is colonized. Note that we do not always require that both single-drug compartments exist, and the compartment sizes may not follow this particular geometric relationship.

To describe population dynamics of the pathogen in this scenario, we use a viral dynamics model(31) (Suppl. Methods) which tracks infected and uninfected cells. We analyze the model using a fully stochastic simulation (see Suppl. Methods), and derive approximate analytic formulas to describe the dominant processes. The exact model choice is unimportant, as any model of pathogen growth with limiting resources, such as the logistic model, will give nearly identical results. In this model, pathogen fitness can be measured in terms of the *basic reproductive ratio* R_0_, the number of new infections generated by a single infected cell before it dies, when target cells are in excess. A strain can only lead to a sustainable infection in a compartment if R_0_ > 1 (i.e., growth is positive). When this occurs, the pathogen population can reach an equilibrium level which we refer to as the *carrying capacity* (*K*).

We consider at most four compartments within a single patient (Figure 1): one compartment where no drugs are present (the *sanctuary*), two compartments where only one of the drugs is present (*single-drug compartments* 1 and 2) and one compartment where both drugs are present (the *double-drug compartment*, which we always take to be by far the largest compartment). The pathogen population within each compartment is assumed to be well-mixed and follows the viral dynamics model. The size of each compartment *j* is given by the number of target cells *N*_*j*_ that it contains when infection is absent. The carrying capacity *K*_*ij*_ of pathogen strain *i* infecting compartment *j* increases monotonically with pathogen fitness 
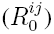
 and is always less than the compartment size (*K*_*ij*_ < *N*_*j*_ for all *i*), assuming that the death rate of infected cells exceeds that of uninfected cells. In the absence of mutation or migration, there is competitive exclusion between strains within a compartment, and the strain with the highest fitness goes to fixation. With migration or mutation, multiple strains may co-exist within a compartment, although the locally suboptimal strains generally occur at much lower frequencies.

The four compartments are connected by migration of pathogens (but not drugs), and every strain in the body migrates from compartment *j* to compartment *k* at a rate *m*_*jk*_. We use a simple and biologically realistic migration scheme in which each pathogen migrates out of its home compartment at the same rate *m*. Migrants from a given compartment are then distributed into all four compartments (including the one they came from) proportionally to the compartment sizes, so that larger compartments get more migrants.

A single mutation is needed for resistance to each drug. Mutations conferring resistance to drug *i* occur at a rate *μ*_*i*_ (and can revert at the same rate). Resistance to two drugs requires that both mutations occur, and we do not allow recombination to occur. We design our fitness landscape so that four assumptions are met: i) Each drug alone suppresses pathogen growth, ii) A wild-type pathogen in the sanctuary has the highest possible fitness, iii) A doubly-resistant pathogen is always viable, and iv) In the single-drug compartments, the strain with resistance only to the drug present is the fittest. Formally, if a strain is *not* resistant to a drug *i* present in the compartment where it resides, its fitness is reduced by a factor of 1 − ϵ_*i*_, where ϵ_*i*_ ∈ [0, 1] is the drug efficacy. Resistance mutations come with a fitness cost *s*_*i*_ ∈ [0, 1]. The fitness of resistant strains is completely unaffected by the presence of the drug. To satisfy condition (i), we constrain *R*_*wt*_(1 − ϵ_*i*_) < 1, and to meet condition (iii), we require *R*_*wt*_(1 − *s*_1_) (1 − *s*_2_) > 1. The fitness values for each genotype in each compartment, relative to that of a wild-type strain in the sanctuary (*R*_*wt*_), are shown in Table 1. At the start of treatment, we suppose the wild-type pathogen to be present in all compartments; we first focus on the case without pre-existing resistance mutations and later consider how pre-existing resistance alters results.

We apply this model to a physiologic scenario where the double-drug compartment occupies the vast majority of the body and where isolated infections within the small sanctuary or single-drug sites are not life-threatening on their own. Therefore, treatment failure is said to occur when the multi-drug-resistant mutant colonizes the double-drug compartment. We investigate how the presence and size of single-drug compartments – created by combinations of drugs with mismatched penetration profiles – determine two clinical outcomes: the rate at which treatment failure occurs, and the evolutionary path by which the multi-drug resistant mutant emerges. Under the *direct evolutionary path*, multiple resistance mutations are acquired near-simultaneously (this is sometimes referred to as “stochastic tunnelling” (32, 33)); under *stepwise evolution*, a single-drug compartment is colonized with a single-resistant strain prior to the emergence of multi-drug resistance.

**Table 1:**
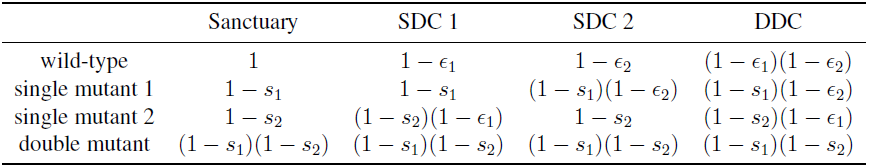
The fitness of each pathogen strain in each compartment relative to the fitness of the wild-type strain in the absence of the drug (*R_wt_*). The efficacy of drug *i* is ϵ*_i_* and the fitness cost of resistance to drug *i* is *s_i_*. SDC: single-drug compartment. DDC: double-drug compartment.

## 3 Results

### 3.1 Mismatched drug penetration can speed up emergence of resistance

Using parameter values appropriate for HIV treatment (see Supplementary Methods), we simulate pathogen evolution according to the model described above. For simplicity, we first consider the presence of only one single-drug compartment (containing drug 1). The probability of treatment failure via double-drug resistance after one year (Figure 2a) or 10 years (Fig 2b) increases dramatically with the size of single-drug compartment, even when this region is 2 – 3 orders of magnitude smaller than the area covered by both drugs. This demonstrates that imperfect drug penetration can be highly detrimental to treatment outcomes.

Mismatched drug penetration hastens the emergence of multi-drug resistance by allowing for stepwise evolution (Fig 2c, d). Specifically, single-resistant mutants can evade competition with wild-type strains by migrating to the single-drug compartment, which serves as a platform from which resistance to the second drug may evolve (Fig 2d). When drugs have identical penetration, there are only two compartments – the sanctuary and the double-drug compartment. In typical simulations (Fig 2c), single-resistant mutants arising in the sanctuary are driven recurrently to extinction by the fitter wild type. As a result, the only way that double-drug resistance can emerge is by appearance of both mutations nearly simultaneously, enabling successful migration to the double-drug compartment (direct evolution). This slow process increases time to treatment failure.

Consistent with the above explanation, the prevalence of failure by direct evolution depends weakly on single-drug compartment size, only decreasing slightly with compartment size as the competing stepwise path occurs first (Fig 2a, b). Failure by stepwise evolution, however, increases substantially with the size of the single-drug compartment, and it is the dominant path if the single-drug compartment exceeds a critical size, investigated below.

**Figure 2:**
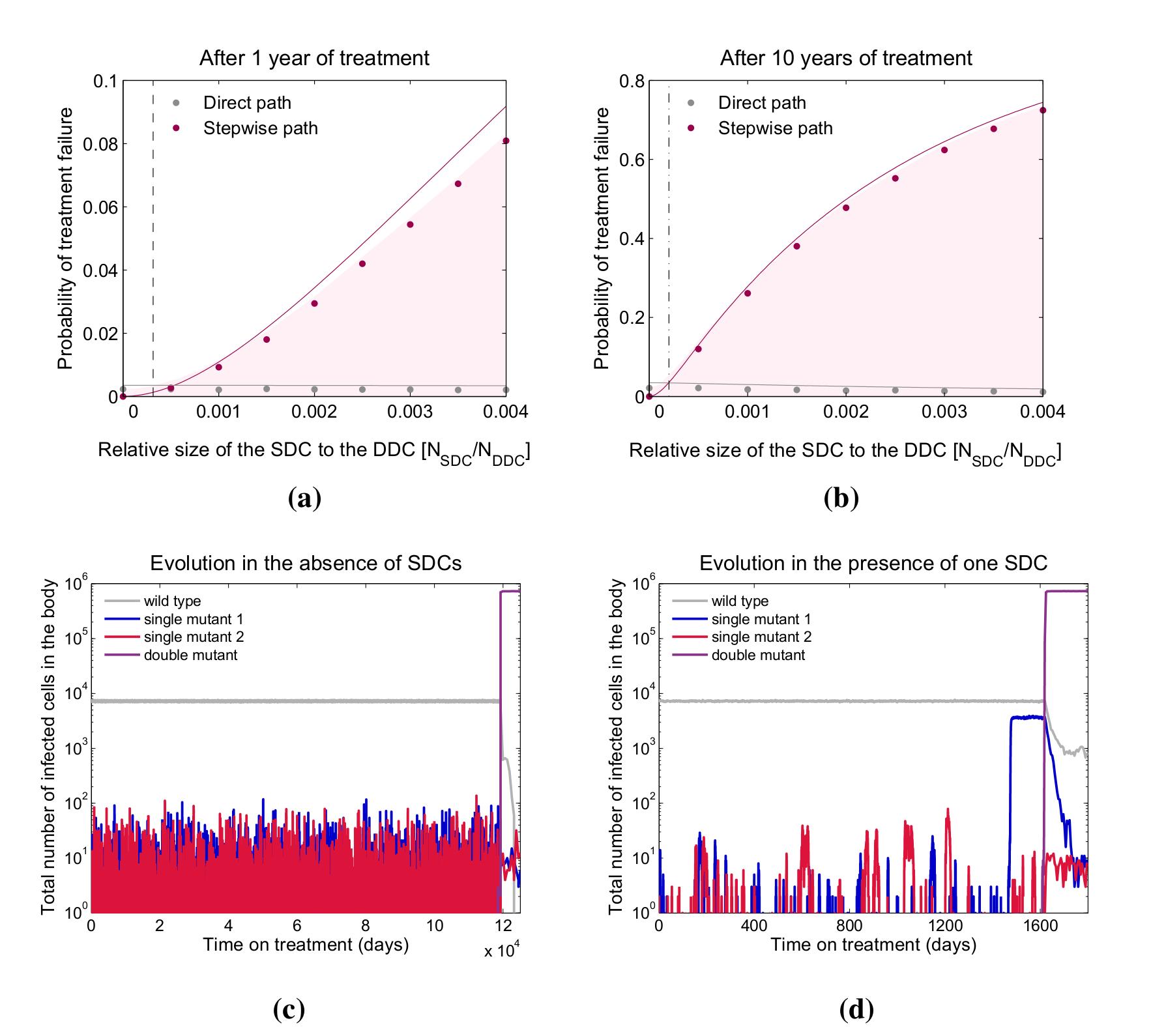
Resistance evolution in the presence of a single-drug compartment. Even a small single-drug compartment can considerably speed up the evolution of double drug resistance. a-b) The shaded area gives the fraction of simulated patients that failed treatment after 1 or 10 years as a function of the size of the single-drug compartment containing drug 1 (SDC1) relative to the size of the double-drug compartment (DDC). We further indicate whether treatment failure occurred via direct (grey dots) or stepwise evolution (pink dots). Solid lines are analytic calculations (Suppl. Methods §5, 6). The vertical dotted lines are analytical approximations for the point where the stepwise path to resistance becomes more important than the direct path (§3.2, Supp. Methods §7). c-d) Evolution of drug resistance over time for a simulated patient in the absence (c) or presence (d) of SDC1. When there are no single-drug compartments, mutants resistant to drug 1 go to extinction recurrently by competition with the wild type in the sanctuary, whereas in the presence of SDC1, mutants resistant to drug 1 can escape competition and establish a continuous population (blue line) from which a doubly-resistant strain can evolve (purple). Parameters: *R*_*wt*_ = 4; _1_ = 0:99; _2_ =0:99; *d*_*y*_ = 1; *d*_*x*_ = 0:1; *m* = 0:1; *s*_1_ = 0:05; *s*_2_ = 0:05; _1_ = 10 ^5^; _2_ = 10 ^5^; *N*_*SAN*_ = 10^5^; *N*_*SDC*2_ = 0; *N*_*DDC*_ = 10^7^. *N*_*SDC*1_ changes along the x-axis for a) and b) and for each value of *N*_*SDC*1_ treatment has failed in at least 1000 simulated patients. *N*_*SDC*1_ = 0 for c) and *N*_*SDC*1_ = 5 × 10^4^ for d).

### 3.2 Stepwise versus direct evolution

Using a simplified model of colonization of each compartment, we can approximate the critical single-drug compartment size above which stepwise evolution becomes the dominant process. Specifically, we approximate the colonization process by transitions between discrete states of the population, where each state is described by the presence or absence of each strain in each compartment. For brevity, we assume that the mutation rate and fitness cost are the same for both mutational steps and that there is only one single-drug compartment. In State 0, only the sanctuary is colonized (by the wild-type strain); in State 1, the single-drug compartment is also colonized (by the single-resistant mutant); and State 2 is the end-state where the double-drug compartment is colonized (by the double resistant strain). Rates of treatment failure can be computed exactly in this simplified model (Suppl. Methods §4-6), which provides an excellent approximation to the full stochastic simulation (Figure 2a, b).

Using this model, we can obtain simple approximate expressions for the size of the single-drug compartment (SDC) where the stepwise path starts to overtake the direct path (detailed in Suppl. Material §7). The SDC becomes colonized (transition from State 0 to 1) by one of two events. Either a mutation occurs within the sanctuary, and then that strain migrates to the SDC, or, a wild-type strain migrates from the sanctuary to the SDC, where it manages to replicate and mutate despite the presence of the drug. In both cases the mutant must escape extinction to establish an infection in the SDC. In the limit where mutation cost is small (s << 1) but drug efficacy is high (ϵ ~ 1), mutation typically precedes migration, and the rate of invasion of the single-drug compartment is approximately

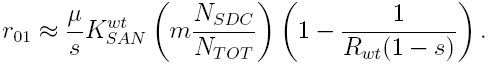

Here 
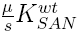
is the number of single mutants in the sanctuary (“SAN”, at mutation-selection equilibrium), 
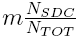
 is the migration rate to the single-drug compartment, and 
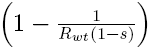
 is the establishment probability of a resistant mutant in the single-drug compartment (see Suppl. Methods for full derivation with respect to viral dynamics model). If invasion is successful, we assume that the population in the newly invaded compartment reaches its carrying capacity 
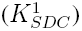
 instantaneously. Doing so relies on a separation of timescales between the slow processes of mutation and migration, and the faster process of growth to equilibrium.

Similarly, once the single-drug compartment is colonized, the double-drug compartment (DDC) can be invaded. Again, the mutation-migration path is most likely, with rate approximately

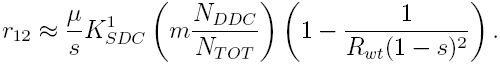

The double-drug compartment can also be invaded directly from the sanctuary. There are three paths by which this can happen, depending on whether none, one, or both of the necessary mutational steps occur before migration. By the same logic as above, the mutation-mutation-migration path is most likely, and the rate is approximately

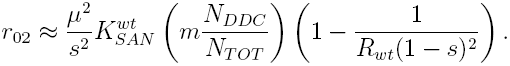

In the scenario under consideration, the mutation rate is much smaller than the cost of mutations 
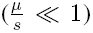
 so that 
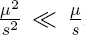
. We also assume that both drugs penetrate in a large part of the body so that the double-drug compartment is always much larger than the single-drug compartment (*N*_*SDC*_ << *N*_*DDC*_). It is therefore likely that *r*_01_ << *r*_12_ and *r*_02_ << *r*_12_. Using these expressions, we can determine the overall rate at which the DDC becomes colonized via the SDC (stepwise evolution), and compare it to the rate of direct evolution.

First, we consider treatment outcomes when a short enough time (*t*) has passed so that drug resistance is rare and all steps are rate-limited (*r*_01_*t* << 1, *r*_12_*t* << 1, *r*_02_*t* << 1). In this regime, the minimum size of the SDC at which stepwise evolution outpaces the direct path (lines cross in Figure 2a, b) increases with the pathogen virulence 
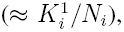
, but decreases with the migration rate (*m*) and (weakly) with the fitness of the single mutant (*R*_*wt*_(1 − *s*)). It also decreases with the time of observation (*t*) because the stepwise path requires two steps, so that for very small t, the SDC needs to be larger for it to be possible that both steps are completed. This approximation (Suppl. Methods §7, Approx. 1) describes the cross point after 1 year of treatment well (Figure 2a). Alternatively, if the treatment time is long enough so that most individuals who developed single drug resistance progressed to treatment failure (*r*_12_*t* > 1), but the other (slower) steps remain rate limiting, then a simpler and more intuitive result emerges: the stepwise path is more important than the direct path if

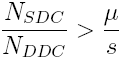

The single-drug compartment therefore plays an important role if its size, relative to the double-drug compartment, is at least equal to the mutation-to-cost ratio. Intuitively, if mutations are rare and costly, then double mutants occur infrequently and the stepwise path to multi-drug resistance is relatively more important. Even if mutations are rather common (say, *μ* = 10^−5^) and not very costly (*s* = 10^−3^), the stepwise path is still dominant if the single-drug compartment is at least one hundredth the size of the double-drug compartment. For the parameters used in the figures, this approximation describes the cross point after 10 years of treatment well (Figure 2b).

### 3.3 Trade-off between halting pathogen growth and preventing resistance

Minimizing the size of single-drug compartments can impede the stepwise evolution of resistance. Pursuing this goal, however, involves choosing drugs that penetrate the same anatomical regions, potentially reducing the portion of the body that receives any drug at all. The physician is therefore faced with a trade-off: to halt wild-type growth immediately (smaller sanctuary) or to prevent stepwise evolution of resistance (smaller single-drug compartments)?

To investigate this trade-off, we vary single-drug compartment size relative to the sanctuary, keeping double-drug compartment and total system size constant (Fig 3). Consistent with the above findings, the rate of treatment failure by double-drug resistance increases dramatically as single-drug compartment size increases from zero. At the same time, however, the sanctuary shrinks from its maximum size, reducing total pathogen load prior to failure.

The trend in treatment failure reverses, however, as the single-drug compartment grows larger than the sanctuary (right half of Fig 3): although stepwise evolution is increasingly the dominant mode of treatment failure, a small sanctuary limits the rate at which single mutations can occur. In the absence of a sanctuary, treatment failure can occur only if there is a pre-existing resistance mutation or if the pathogen acquires resistance shortly after treatment starts. In this limit, the dynamics are a classic “race to rescue” described by Orr and Unckless(34). The rate of treatment failure is greatest when the sanctuary and single-drug compartment are the same size, highlighting the fact that stepwise evolution is driven by interaction between the two compartments. These findings suggest that eliminating all sanctuary sites should be a primary goal, but if this is not feasible (for example, if the pathogen has a latent phase not targeted by treatment), then preventing zones of single-drug coverage should take precedence.

### 3.4 Accounting for pre-existing mutations

To focus clearly on the processes by which resistance is *acquired* during combination therapy, we have so far ignored the contribution of pre-existing mutants (known in the population genetics literature as standing genetic variation). To instead include this factor, we simulate the model for a period of time prior to the introduction of treatment, allowing both single and double-resistant mutants to occur along with the wild-type strain in each compartment. Previous work has focused extensively on comparing the relative roles of pre-existing and acquired resistance in viral dynamics models (17, 35–37), and here we simply summarize the trends in our model.

The addition of pre-existing resistance acts to increase the overall rate of treatment failure, and this increase is more prominent for certain parameter values and for smaller treatment times (compare Fig S1a with Fig 2a). However, the inclusion of pre-existing resistance does not affect any of the general trends, such as the dominant path to resistance (Fig S1) or the trade-off between the size of the sanctuary and the single-drug compartment (Fig. 3). Importantly, the role of pre-existing resistance - defined as the percent of failures attributable to standing genetic variation - increases dramatically with single-drug compartment size. Therefore, the presence of compartments where only single drugs penetrate can increase the rate of treatment failure both by making it quicker to acquire multiple resistance mutations and by selecting for pre-existing single-drug resistant mutants.

**Figure 3:**
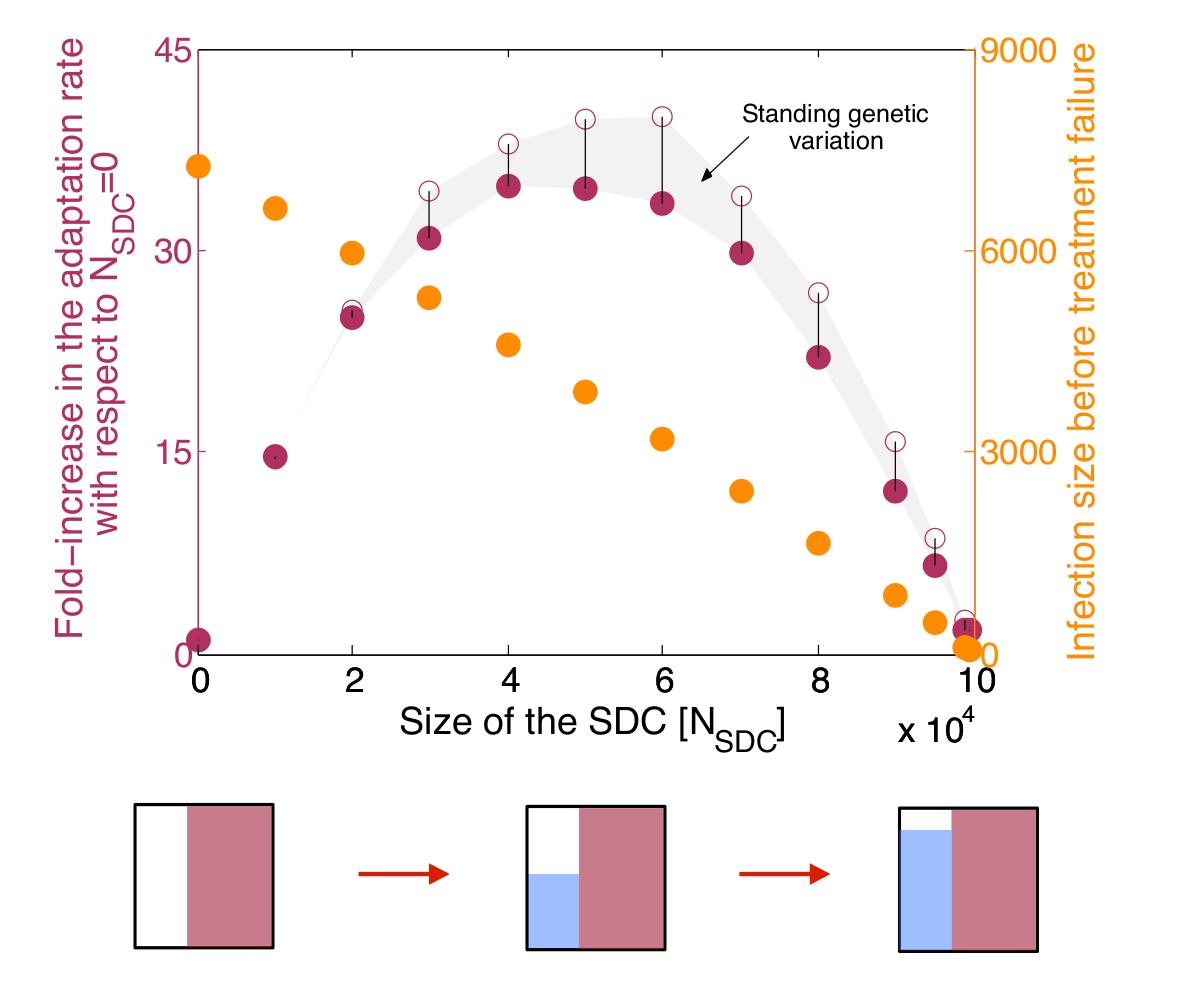
Trade-offbetweentotal drug coverage and the presence of single-drug compartments. The adaptation rate (purple dots, left y-axis) and time-averaged infection size (orange dots, right y-axis) are plotted as a function of the size of the single-drug compartment with drug 1 (*N*_*SDC*1_), assuming that the sum of the sizes of the sanctuary (*N*_*SAN*_) and SDC1 is constant. Diagrams below the x-axis illustrate the changes in compartment sizes, following the style of Figure 1. The adaptation rate is defined as the inverse of the mean time to treatment failure, and is plotted relative to the rate when *N*_*SDC*1_ = 0. We show adaptation rate without (filled dots) and with (open dots) standing genetic variation (i.e., pre-existing resistance); the difference is shown by the grey area and the vertical lines. The infection size is calculated as the mean of the time-averaged number of infected cells in all compartments before treatment failure occurs. Increasing the size of the single-drug compartment provides better control of the infection before treatment fails, but strongly favors resistance evolution if the reduction of the sanctuary is not large enough. The relative contribution of standing genetic variation to failure increases with the size of the SDC. Parameters: *R*_*wt*_ = 4; ϵ_1_ = 0.99; _2_ = 0.99; *d*_*y*_ = 1; *d*_*x*_ = 0.1; *m* = 0.1; *s*_1_ = 0.05; *s*_2_ = 0.05; μ_1_ = 10^−5^; μ_2_ = 10^−5^; *N*_*SAN*_ = 10^5^ − *N*_*SDC*1_; *N*_*SDC*2_ = 0; *N*_*DDC*_ = 10^7^. Each point is an average over 1500 simulated patients.

### 3.5 Order of mutations

Since pharmacological factors determining penetration of anatomical compartments vary widely among drugs (20, 24), we generally expect that each drug in a combination has its own single-drug compartment. In this general case, we can ask: To which drug does the pathogen become resistant first? More precisely, if stepwise evolution occurs, is it likely to be through the path *SAN*→*SDC*1→*DDC* or SAN→SDC2→DDC? Examining the rate of each path as a function of the size of each single-drug compartment (Fig 4a) shows that resistance is more likely to emerge first to the drug with the highest coverage (and therefore largest SDC), and that the odds of resistance occurring to one drug before another are proportional to the ratio of the corresponding SDCs over a large parameter range.

Moreover, the mutation rates and costs associated with resistance to each drug may differ, also influencing the likelihood of a particular path to resistance. Resistance is more likely to emerge first for the drug associated with the highest mutation rate (Figure 4b) and lowest fitness cost (Figure 4c), with the relative rates again being approximated by the ratios of the parameters. Drug efficacy may also vary, though in the regime where each drug individually suppresses wild-type pathogen growth (*R_wt_* (1−ϵ) ≪ 1) and the cost of mutations is not too high (s < ϵ), drug efficacy barely influences the path to resistance.

**Figure 4:**
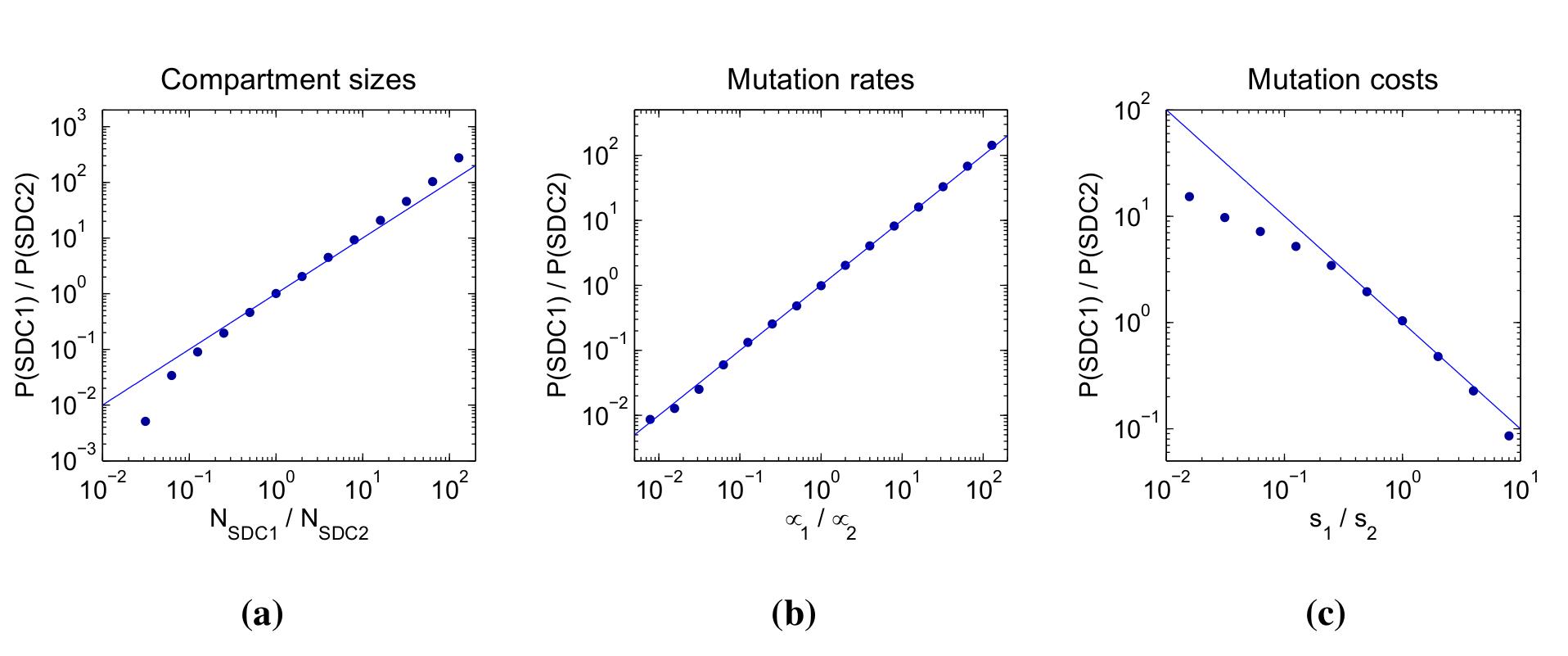
Stepwise resistance evolution in the presence of two single-drug compartments. Fraction of simulated patients that failed via the path where the single-drug compartment with drug 1 is colonized before treatment failure (P(SDC1): *SAN*→*SDC*1→*DDC*) relative to the fraction that failed via the path where the single-drug compartment with drug 2 is colonized before (P(SDC2): *SAN*→*SDC*2→*DDC*) as a function of a) compartment sizes, b) mutation rates and c) mutation costs. a) The x-axis corresponds to the ratio of the size of the single-drug compartment with drug 1 (*N*_*SDC*1_) to the size of the single-drug compartment with drug 2 (*N*_*SDC*2_). b) The x-axis corresponds to the ratio of the mutation rate for resistance to drug 1 (μ_1_) to the mutation rate for resistance to drug 2 (μ_2_). c) The x-axis corresponds to the ratio of the cost of a resistance mutation to drug 1 (*s*_1_) to the cost of a resistance mutation to drug 2 (*s*_2_). Simulation results (dots) are overlaid with the lines *y* = *x* (for a and b) or *y* = 1/*x* (for c). Parameters: *R*_*wt*_ = 4; ϵ_1_ = 0.99; ϵ_2_ = 0.99; *d*_*y*_ = 1; *d*_*x*_ = 0.1; *m* = 0.1; *s*_1_ = 0.05; *s*_2_ = 0.05; μ_1_ = 10^−5^; μ_2_ = 10^−5^; *N*_*SAN*_ = 10^−5^; *N*_*SDC*1_ = 1 × 10^4^; *N*_*SDC*2_ = 1 × 10^4^; *N*_*DDC*_ = 10^7^. *N*_*SDC*1_ changes along the x-axis for Figure 4a, μ_1_ changes along the x-axis for Figure 4b and *s*_1_ changes along the x-axis for Figure 4c. The total number of simulated patients for each point is 3000.

## 4 Discussion

Antimicrobial drugs fail to reach effective concentrations in many tissues and body organs, allowing pathogen replication and potential evolution of resistance (22, 25, 38). We studied the role of imperfect drug penetration in the development of drug resistance during combination therapy using a model of within-host pathogen evolution. In particular, we focused on the consequences of mismatched drug penetration, which may be common during combination therapy (18, 20). Our findings are summarized in Figure 5.

**Figure 5:**
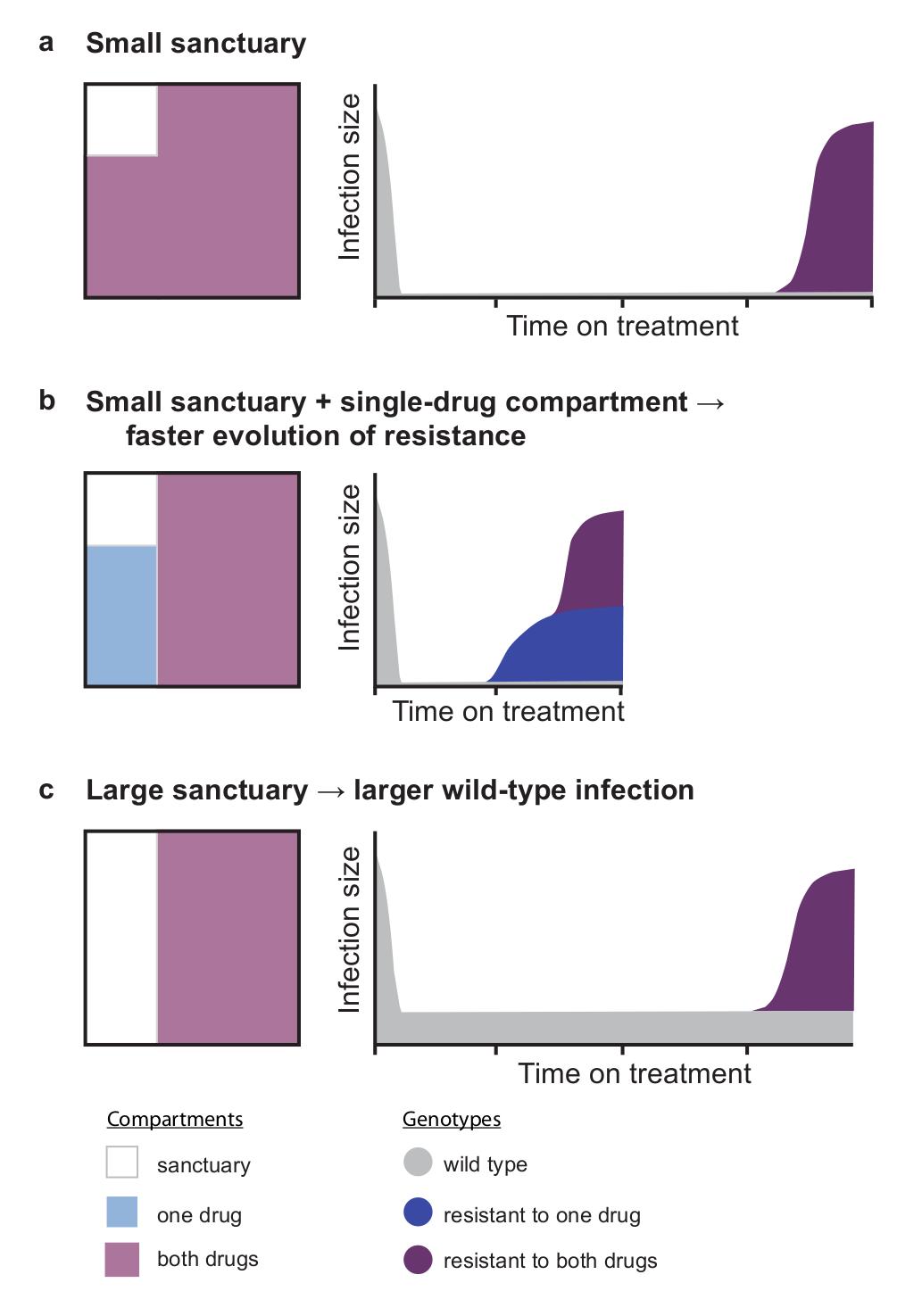
Summary of the evolution of resistance with imperfect drug coverage. a) When both drugs have high, matched penetration throughout the body, the evolution of multi-drug resistance is slow, since it requires either pre-existing multi-drug resistance or near-simultaneous acquisition of both mutations along with migration out of a sanctuary site. If one drug (b) or both drugs (c) have a lower penetration, treatment outcomes may suffer in different ways. b) If there are regions where only one drug reaches an effective concentration, then the evolution of multi-drug resistance speeds up, because mutations may emerge in a stepwise fashion via single-drug compartments. Single mutations can arise *de novo* from wild-type pathogen in the sanctuary or be selected from pre-existing mutations in the single-drug compartment when treatment is started. c) If the sanctuary is larger but both drugs reach the same regions of the body, then resistance still evolves slowly, but the infection size before treatment failure will be larger. Therefore if high penetration of all drugs is impossible, there is a trade-off when choosing which drugs to pair in combinations: halting growth of the wild-type pathogen immediately (b), or preventing the sequential accumulation of resistance mutations (c).

In this model, mismatched penetration of two drugs into anatomical compartments sped up the evolution of multi-drug resistance dramatically by creating zones of spatial monotherapy where only one drug from a combination regime is at therapeutic concentration. These zones, or “single-drug compartments,” positively select for single-drug resistant mutants, thereby favoring the fast stepwise accumulation of resistance mutations (Figure 2a, b, d, 5b). Stepwise resistance evolution is hindered when drugs have identical penetration profiles, because in that case single-drug resistant mutants compete with the fitter wild type in the sanctuary and they therefore suffer recurrent extinction (Figure 2c, 5a, c). Without access to the stepwise path, resistance mutations must be acquired near-simultaneously; the system thus takes a far slower “direct” path to treatment failure. Even slight differences in penetration of co-administered drugs lead to a high risk of multi-drug resistance, since the stepwise path dominates the direct path even for very small single-drug compartments (Figures 2a and b).

Although mismatched drug penetration favors resistance evolution, it may sometimes be advantageous to select a combination of drugs with different penetration profiles, as doing so can increase total drug coverage in the body. Such a combination may not only control disease symptoms associated with total pathogen load, but it may also slow the evolution of resistance if it eliminates most sanctuary zones in the body (Figure 3). If elimination of sanctuaries is not possible, however, then avoiding mismatched penetration in a combination regime should be the main strategy for preventing multi-drug resistance. Previous work has questioned the orthodoxy that “aggressive” antimicrobial chemotherapy is optimal for preventing resistance(39, 40). If we consider that one aspect of treatment aggression is the extent of drug penetration, then our model demonstrates the complexities involved in answering this question.

In particular, our model offers an explanation for why the strategy suggested for some antibiotic treatments of pairing a broadly-penetrating drug (e.g., rifampicin) with a narrowly-penetrating one (e.g., vancomycin) to increase total drug coverage (41) might fail frequently due to the rapid evolution of resistance against the drug with higher penetration (41–43). It also offers an alternative explanation to why certain drugs are more vulnerable to resistance. This vulnerability is usually explained by a low genetic barrier to resistance (i.e., only one mutation needed) or by their long half-life. Our model suggests that broad penetration may also make a drug vulnerable to the evolution of resistance, if the drug is paired with drugs with lower penetration (Figure 4).

Our model also offers an explanation of stepwise evolution of resistance in HIV infection, the commonly observed pattern whereby the virus gains one resistance mutation at a time (15, 44, 45). As treatment regimes are designed so that each drug is active against mutants resistant to the others, single-resistant mutants should be driven to extinction both in sanctuary zones (by competition with fitter wild type) and where all drugs are active (by sensitivity to all drugs save one). It has been hypothesized that either non-adherence to treatment or different drug half-lives cause effective *temporal* monotherapy, which is to blame for the appearance of single-drug resistant viruses(13, 14). We propose that mismatched penetration of drugs in a combination treatment offers an alternative explanation for this stepwise evolution of resistance, via effective *spatial* monotherapy. Very small single-drug compartments are sufficient to cause this effect, suggesting that these regions may be very hard to detect and could remain overlooked.

In this study, we focused on the case of treatment with two drugs, but we expect that our results generalize to three or more different drugs. Adding a third drug to a regimen may reduce the size of the sanctuaries and/or the size of single drug compartments and should therefore reduce the rate at which multi-drug resistance evolves.

Several extensions to our model can be considered in future studies. Firstly, we assume that drug compartments are discrete and have a fixed size, however, drug concentrations can be continuous in space and the pharmacokinetics of individual drugs can modify the size of the different compartments over time. Secondly, we have assumed that treatment fails when the double-drug compartment is invaded, but depending on the size and location of drug compartments in the body, treatment may fail when a single-drug compartment is invaded. Also, we assume a very specific migration model between the compartments (known in population genetics as the island model, (46, 47)) but other migration models may be possible. Specifically, not all compartments may be connected by migration and the migration rates may be independent of the size of the target compartment.

Drug compartments are commonly described as specific anatomical locations in the body like organs or tissues. For instance, not all antimicrobial drugs penetrate to therapeutic concentrations in the central nervous system (18, 21, 22), the genital tract (18, 20), and the lymphoid tissue (18, 38). However, the compartments in our model could be interpreted in many ways. For example, they could represent different cell types, such as phenotypically resistant subpopulations of bacteria that have low permeability to antibiotics (48) or cells in a tumour that are not reached by anticancer drugs (49, 50). Compartments could also exist at a population level, representing differential targeting of geographic regions with insecticides, herbicides or therapeutics. Finally, this model might be relevant to other evolutionary processes where multiple adaptations are ultimately needed for survival and to the study of the role of spatial heterogeneity on adaptation.

## 5 Materials and Methods

We use a basic viral dynamics model (31) to simulate the infection within each compartment and we include stochastic mutation and stochastic migration among all the compartments. We perform exact stochastic simulations, tracking the genotype and location of every infected cell in the body and explicitly simulating all the events that might occur to a cell: replication (representing either division of a bacterial cell or infection of a new cell by a virus), mutation (upon replication), death and migration among different compartments. Simulations are performed using the Gillespie algorithm. Details of the model, analytic approximations, and simulation methods are provided in the Supplemental Methods.

## 6 Acknowledgments

The authors thank A. Harpak, P. Altrock, J. Wakeley, J. Hermisson, D. Weissman, H. Uecker, H. Alexander, B. Waclaw, S. van Doorn and P. Abel zur Wiesch for helpful feedback on the project and manuscript at various stages. S. M. G was funded by the Erasmus Mundus Masters Programme in Evolutionary Biology. A. L. H. and D. I. S. R. were supported by Bill and Melinda Gates Foundation Grand Challenges Explorations Grant OPP1044503. A. L. H was also supported by National Institutes of Health grant DP5OD019851. D. A. P was supported by National Institutes of Health grants RO1GM100366 and RO1GM097415. M. A. N. was supported by the John Templeton Foundation. P. S. P. received funding from the Human Frontiers Science Program (LT000591/2010-L)

